# System identification reveals multiple interacting states in visuomotor adaptation

**DOI:** 10.1101/2022.07.11.498664

**Authors:** Priscilla Balestrucci, Marc Ernst

## Abstract

Many characteristics of sensorimotor adaptation are well predicted by the Kalman filter, a relatively simple learning algorithm for the optimal estimation of dynamic variables given noisy measurements. Yet not all Kalman filter predictions are confirmed empirically, suggesting that the model might not be sufficient to describe the observed behavior. In our study, we propose that sensorimotor adaptation can be modeled with multiple interacting states, each one described as a Kalman filter, to better reflect the architecture plant of the physical system implementing the behavior (i.e., the different motor and sensory components involved in adaptation). To test our hypothesis, we measured motor error in a series of rapid reaching tasks in which we introduced different conditions of feedback uncertainty and systematic perturbations. We then applied system identification procedures to the resulting adaptation response to test which system architecture would best fit the data. We considered three possible architectures: one with a single Kalman filter, with two filters in series, or two filters in parallel. We found that this latter structure consistently provided a significantly better fit than the others. When evaluating the identified system parameters, we found that the learning rates of both states decreased under higher uncertainty, while the weight assigned to the slower state increased. Moreover, the dual-state parallel system accounted for the presence of a constant residual bias in the adaptation to a step offset, which has been repeatedly reported but is in disaccord with the predictions of the single Kalman filter. We propose that the identified system architecture reflects how the error is assigned to different components of the physical plant responsible for adaptation: namely, as uncertainty increases, the controller assigns a larger contribution in error reduction to the slower state, which is more likely to have been responsible for the measured error.

## Introduction

We constantly adjust and recalibrate the motor plan of our actions over subsequent attempts, based on the difference between desired and perceived outcome. This error correction process is referred to as sensorimotor adaptation, and it is necessary to maintain adequate levels of performance despite changes in the environment, such as those occurring when adjusting to the characteristics of a new tool, and to our own body, for example because of muscle fatigue or body transformations during development (1–3). Short-term adaptation allows us to maintain a highly flexible control of generated movements in different conditions, and it is a strategy used by the nervous system in all kinds of movements, e.g. arm reaching (4), walking (5), eye saccades (6), or speech production (7,8).

Typically, experimental paradigms used to investigate properties of sensorimotor adaptation of arm reaching and pointing movements—which are the focus of the current study—consist in measuring the response to an applied perturbation. Experimental perturbations can consist of a change of the visual feedback displacement associated with the performed movement (9–12), or of the alteration of the motion dynamics obtained by applying a force field via a robotic effector (13–15). When a perturbation is introduced, adaptation is usually fairly rapid, but not instantaneous, as error is reduced incrementally over different trials and not in one single attempt. The fraction of the error corrected from one trial to the next, i.e. the rate of error reduction, is a measure often used to describe and model sensorimotor adaptation (16). Several studies showed that adaptation rate is not constant, instead it depends on many different factors, such as the reliability of the error signal (10,17), the rate of change of environmental statistics (9,10,18), and prior experience with a given perturbation (19–22). Other factors do not seem to have an effect on adaptation rate, in particular movement or perturbation variability (10,23).

Different computational approaches have been applied to describe and predict changes in the adaptation rate. The mathematical formalization used for modeling dynamic, linear time-invariant (LTI) systems is widely used in studies investigating the properties of adaptation from the point of view of motor control (24). In the state-space representation of LTI systems, the input is the sensory error signal induced by the mapping perturbation, the output is the motor adaptation signal, and the hidden states correspond to the estimation of the change that occured to the mapping. While the state-space formalization is very effective in capturing many characteristics of sensorimotor adaptation (21,25,26), it is primarily a descriptive approach, which does not contribute to providing answers to fundamental questions about the characteristics of adaptation (16). In particular, we are interested in understanding, *why* does adaptation rate change instead of being a fixed parameter for an individual? Even more fundamentally, why should there be a rate of adaptation, and error is not completely removed as soon as it is encountered? By adopting a normative rather than descriptive perspective, Bayesian models provide a framework which is better suited to explore such aspects of adaptation. Bayesian models build on the premise that the parameters of adaptation are set and change optimally, based on the knowledge of the statistics of the environment and the uncertainty associated with sensorimotor information (27–29). Since adaptation requires tracking dynamic states and updating the estimation over time given noisy observations, for comparing human behavior to that of an optimal observer, typically a Kalman filter algorithm is applied to model empirical data (9,10,30). In several cases, the predictions of the Kalman filter were confirmed by the experiments. However, not all predictions were validated when comparing human and optimal observer behavior. For example, the Kalman filter predicts that when a constant perturbation is introduced, the error decreases exponentially until it is fully compensated. Instead, several studies showed that adaptation to a constant perturbation is incomplete (10,31,32). Furthermore, movement variability and adaptation rate should be correlated according to the algorithm’s prediction, but such a relation has not been found empirically (10,23).

Despite the fundamental differences between descriptive approaches such as state-space models and normative approaches such as the Kalman filter, their formalizations are nearly identical (2). Using the state-space representation, the Kalman filter can in fact be represented as a dynamic system with a single state, fully described by one free parameter (i.e. the Kalman gain K) which varies according to the statistics of the environment (Fig. 1A). However, given the complexity of the processes underlying sensorimotor adaptation, it is possible that such a model with only a single free parameter is too simple for predicting the behaviors implemented by the physical plant, i.e. the body, responsible for adaptation. Therefore, it is likely that a more articulated representation of the components of the system can instead provide predictions that are better confirmed by empirical data. The idea that multiple free parameters are necessary to describe the characteristics of sensorimotor adaptation has been proposed and successfully validated in multiple studies using the state-space model formalization e.g. (33,34). Smith and colleagues (21) first proposed the existence of at least two interacting adaptive processes, each one operating at different timescales, which could capture several complex adaptation phenomena such as savings (i.e. the faster adaptation occurring when a perturbation is experienced a second time) (35), anterograde interference (the slower adaptation occurring after adapting to a different perturbation) (36), rapid unlearning (extinction of the adaptation effect once the perturbation is removed) (37). Typically, models including multiple interacting states assume that each state is characterized by different learning and forgetting rates, but no hypotheses are proposed concerning the origin of such parameters.

**Figure 1:**
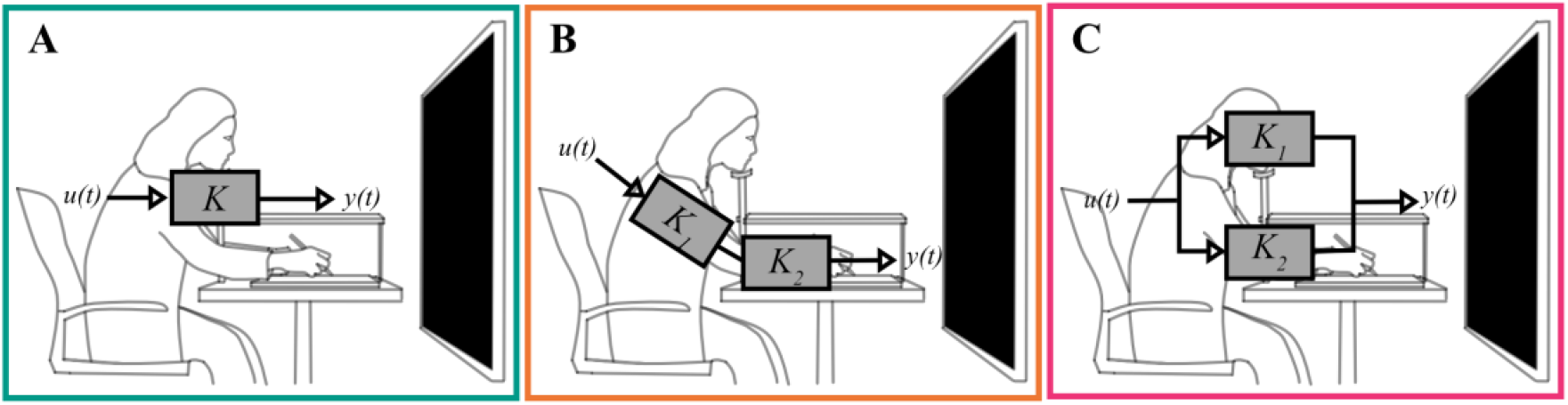
Possible system architectures and relative biological plant. **A)** Single-state system. There is a single stage of adaptation with rate *K*. **B)** Dual-state serial system. Adaptation is the result of two independent states arranged in series, possibly representing the motor system adapting at different joints. **C)** Dual-state parallel system. Two independent states are arranged in parallel, possibly representing different sensory and motor components.

In our study, we hypothesize that the biological plant responsible for adaptation can be modeled with a linear dynamic system composed of multiple interacting states, where each state is described as a Kalman filter. According to this view, the measured adaptation response would depend on changes in the statistics of the environment, as well as on the physical organization of the system itself. In particular, we considered two possible configurations including two independent states: one with a serial architecture, the other with a parallel architecture. In principle, adaptation could be accomplished by implementing either type of system. On the one hand, a system with two (or more) states arranged in series could stand for the articulated motor system adapting at different joints serially aligned along the motor plant (e.g. the shoulder and elbow joints of the arm, Fig. 1B). On the other hand, a system with states operating in parallel would represent different sensory and motor components of the total plant (Fig. 1C), each adapting at its own rate, such as the contribution to the error of the visual system on the one side and the proprioceptive/motor system of the arm on the other. Obviously when introducing multiple adapting states, a possible interaction between them emerges.

In order to test our hypothesis, we measured adaptation in fast reaching movements under different conditions of uncertainty of the visual feedback location. Uncertainty was modulated by applying Gaussian blurs with different standard deviations to the visual feedback. Next, we applied system identification algorithms to assess which of the system architectures introduced above provided the best fit of the experimental data. In order to measure the adaptation response to systematic perturbations of the mapping between motor reach and endpoint location of the feedback, following the method introduced by Hudson and Landy (38), we applied sinusoidal perturbations with different frequencies instead of an impulse-like step-offset perturbation. In their study, they demonstrated the advantage of using sinusoidal perturbations over the more commonly used step-function: Using both empirical and simulated data, they showed that the adaptive response associated with sinusoidal perturbations provide a better signal-to-noise ratio compared to that from step offsets, already with small perturbation amplitudes. As it is possible to use sinusoidal offset functions with small amplitudes, and the change in mapping occurs gradually from one trial to the next, perturbations are less likely to be overtly detected, allowing for less reliance on cognitive strategies to correct errors when participants perform the task. Moreover, while in the case of adaptation to a step function it is necessary to fit an exponential function to the response, we can apply linearized methods for analyzing the response to a sinusoid, thus allowing us to easily use a parametric model to fit the response. In our application of the proposed method, introducing sinusoidal perturbations with multiple frequencies provided an additional advantage: Namely, for each level of feedback uncertainty, we could construct the response of the system in the frequency domain by putting together the parameters extracted from the adaptation signals associated to the different sinusoidal inputs. The response of a system in the frequency domain is mathematically equivalent to that in the time domain (i.e. the impulse response obtained when measuring the adaptation to as step offset) (39). However, by using the former, we could rely on the parameters extracted from multiple experimental runs (one for each sinusoidal frequency) for constructing the overall system response.

### Model Architectures and Simulations

In order to model the different system architectures under consideration for adaptation, we consider the case provided in the experimental task, in which in every trial *t* a systematic perturbation *P*_*t*_ is applied to the endpoint visual feedback (*F*_*t*_) of the reaching hand movement towards a displayed target (see Fig. 5C):

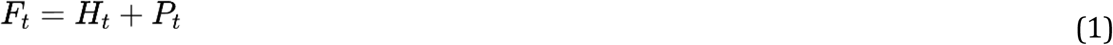

The error on trial *t* results from the distance between feedback and target:

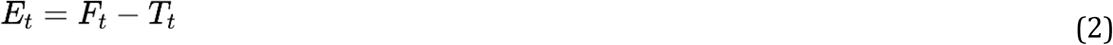

To minimize the error in this task, participants should estimate the visuomotor mapping perturbation based on the performance in previous trials, and make a reaching movement accordingly:

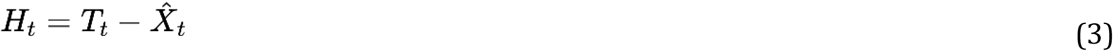

In such conditions, the optimal estimation is provided by the Kalman filter (9,10):

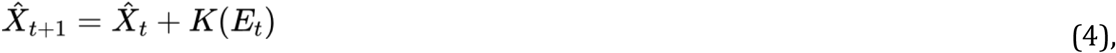

where the Kalman gain *K* defines the speed of the estimation update from one trial to the next, i.e. the rate of adaptation.

Given the identities above (Eq. 1-3), the error can be rewritten as:

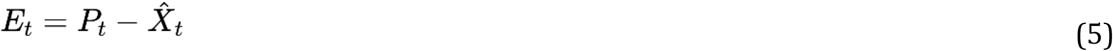

which for the optimal estimation yields:

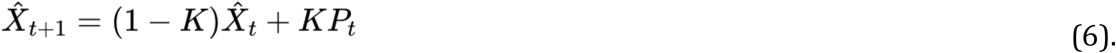

This notation closely resembles the state update of a discrete, LTI state-space system, which is canonically in the form of the difference equation:

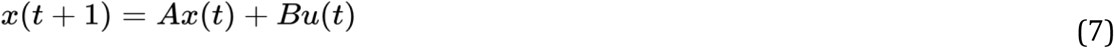

where the input signal *u(t)* is the feedback perturbation *P*_*t*_, and the state update *x(t)* is the mapping estimation 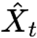.

For the second equation of the state-space system:

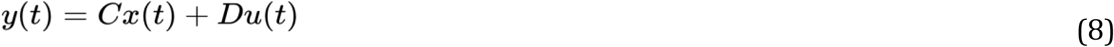

we consider the output of the system as the endpoint error of the hand reaching movement:

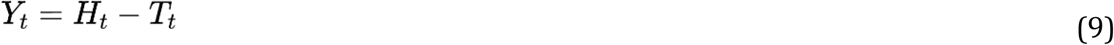

which yields

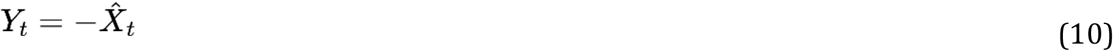

corresponding to the second difference equation of the state-space model where *C = −1, D = 0*.

In the case of a system like the one in Eq. 6, the characteristics of the adaptation response depend on the free parameter of the model, the gain *K*. Such a model resembles a low-pass filter and thus reacts to different input perturbations accordingly: If the input perturbation *P*_*t*_ consists of a constant offset, the model predicts that the adaptation response has the form of an exponential decay with asymptote to 0, while the time constant of the exponential function is given by the Kalman gain *K* (Fig. 2B). For a sinusoidal perturbation, the model predicts that the response has the same frequency, but reduced amplitude and delayed phase with respect to the perturbation. Moreover, the response depends on the free parameter of the model as well as on the speed of the perturbation (i.e. its frequency). The relationship between input and output of a system across frequencies is typically described by the Bode plot of the response in the frequency domain (Fig. 2C). The Bode plot provides the representation of the relative amplitude between input and output expressed in dB (Fig. 2C, top panel) and that of their phase shift in degrees (Fig. 2C, bottom panel).

**Figure 2:**
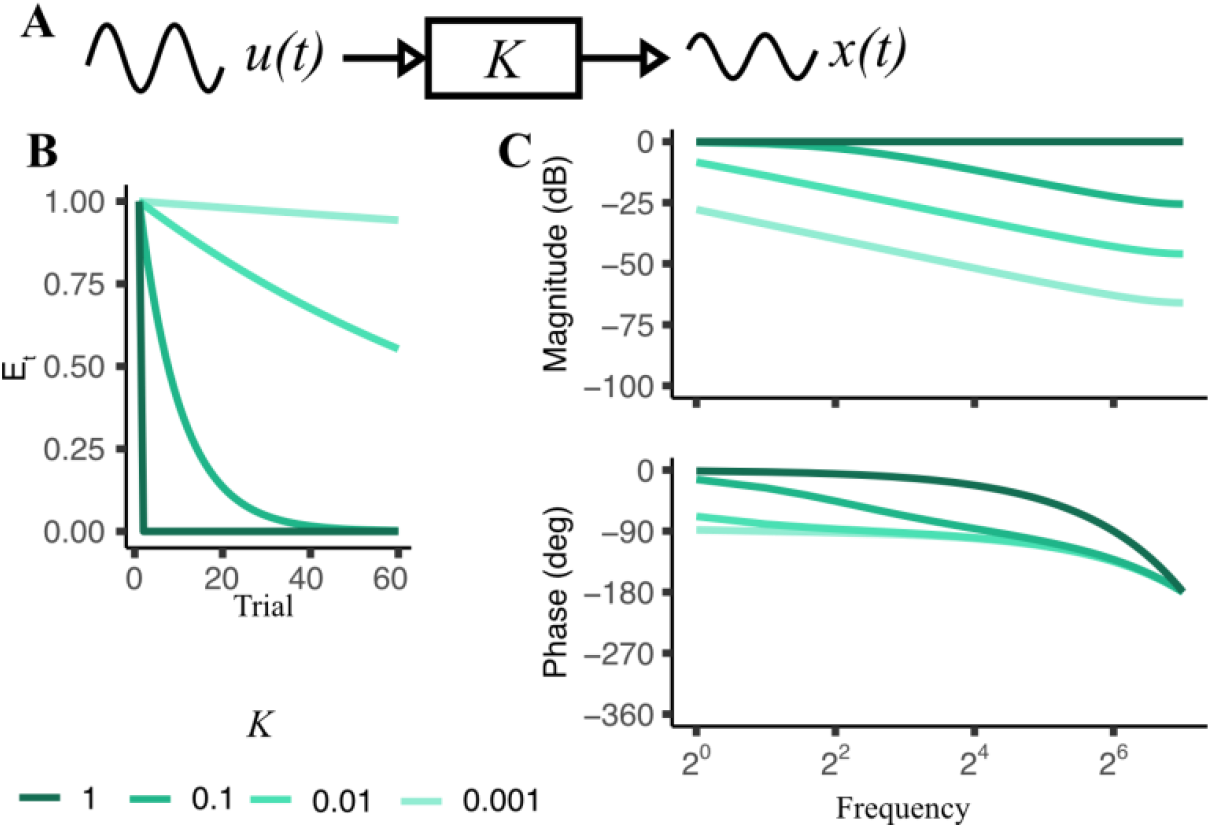
**A)** Single-state system representation. **B)** Response to a step perturbation of a single-state system in the time domain, for varying values of *K*. **C)** Response to sinusoidal perturbations of the same system for varying values of K, in the frequency domain (Bode plot).

Since in our study we measure adaptation to sinusoidal offsets and use such responses to characterize the underlying processes, we will only show simulations of the responses in the frequency domain in the next sections. We used the Matlab’s Control System Toolbox (40) to define the system and to simulate the relative responses in the frequency domain.

### Dual-state systems

In case of a system with two independent states arranged in **series**, the estimation made by the first state provides the input signal for the second, cascading state (Fig. 3A). The estimation updated by the latter, in turn, provides the adaptation output of the overall system. The model is represented by the following system of difference equations:

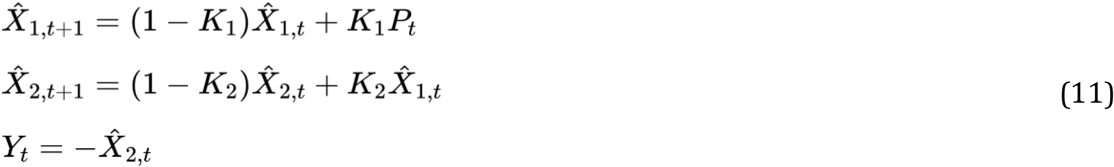

**Figure 3:**
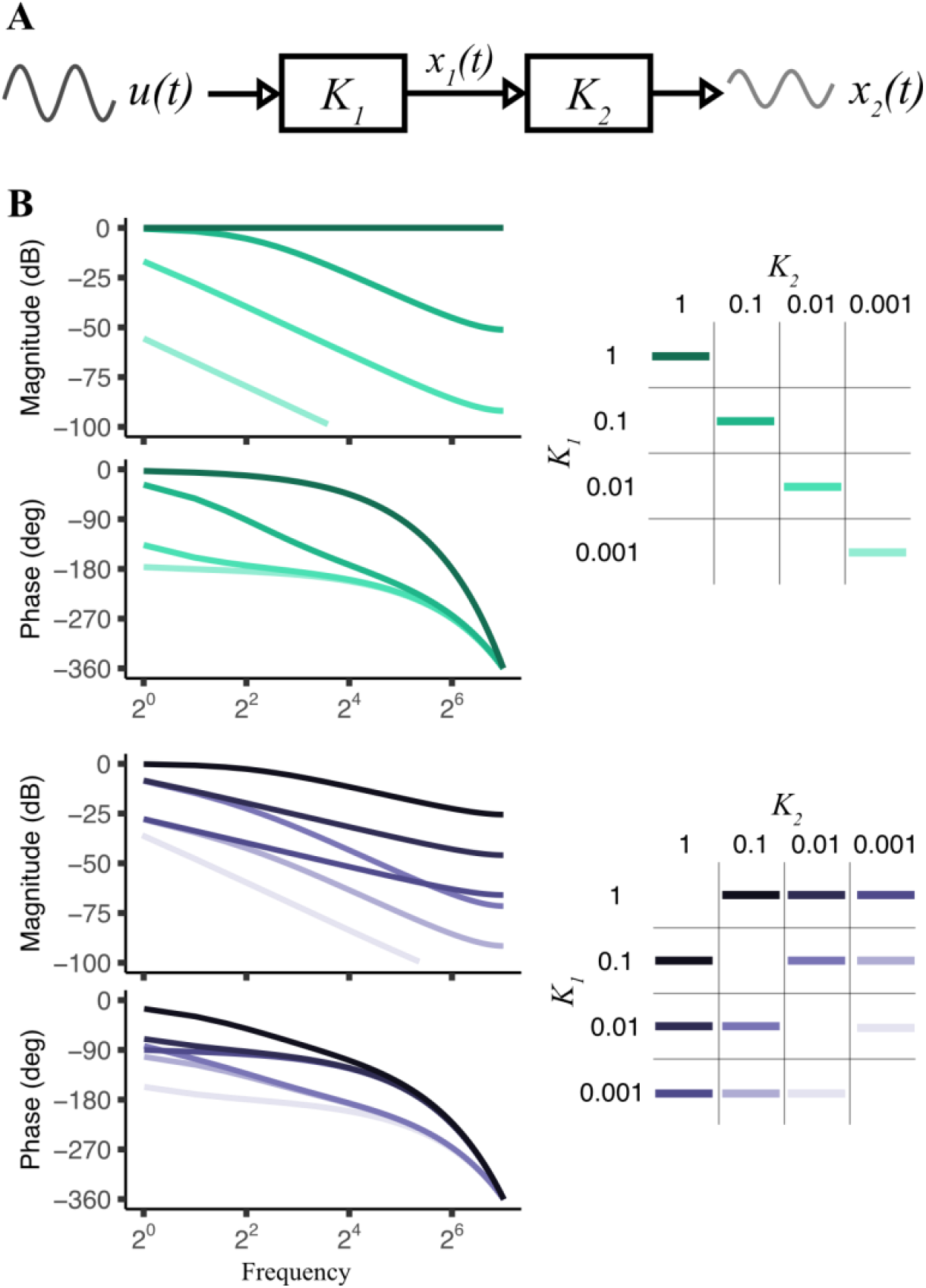
**A)** Representation of a dual-state serial system. **B)** Simulations of the system’s frequency response, for varying values of the two adaptation rates *K*_*1*_ and *K*_*2*_.

The free parameters of the model are the two Kalman gains *K*_*1*_, *K*_*2*_. While the response of such a system is qualitatively similar to that of a single-state system, they are clearly different from each other when considering their frequency responses. In particular, the response of a dual system with two identical states in series is well distinguishable from that of a single state with the same gain (Fig. 2C, 3B-top panel). Moreover, it is not possible to obtain a serial system to a single-state system by setting one of the gains to 0 or 1: In the former case, the response of the serial system is a null response for all frequencies; in the latter, the response of a serial system with gains *K* and 1 differs from that of a single state with gain *K* for a shift in the phase of the response. Note also that the response of the overall system does not depend on the order of each individual state (Fig. 3B, bottom panel).

In a dual system with a **parallel** architecture, each individual state processes part of the input, and the overall output is given by the sum of the estimates performed by each state (Fig. 4A). The system of difference equations describing the model are the following:

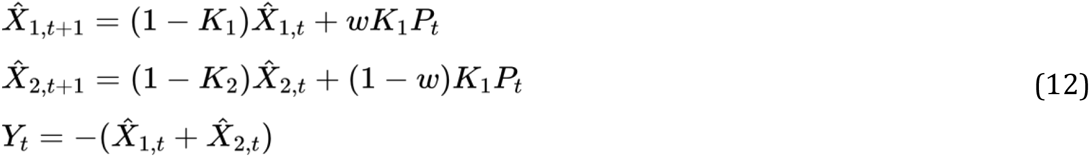

**Figure 4:**
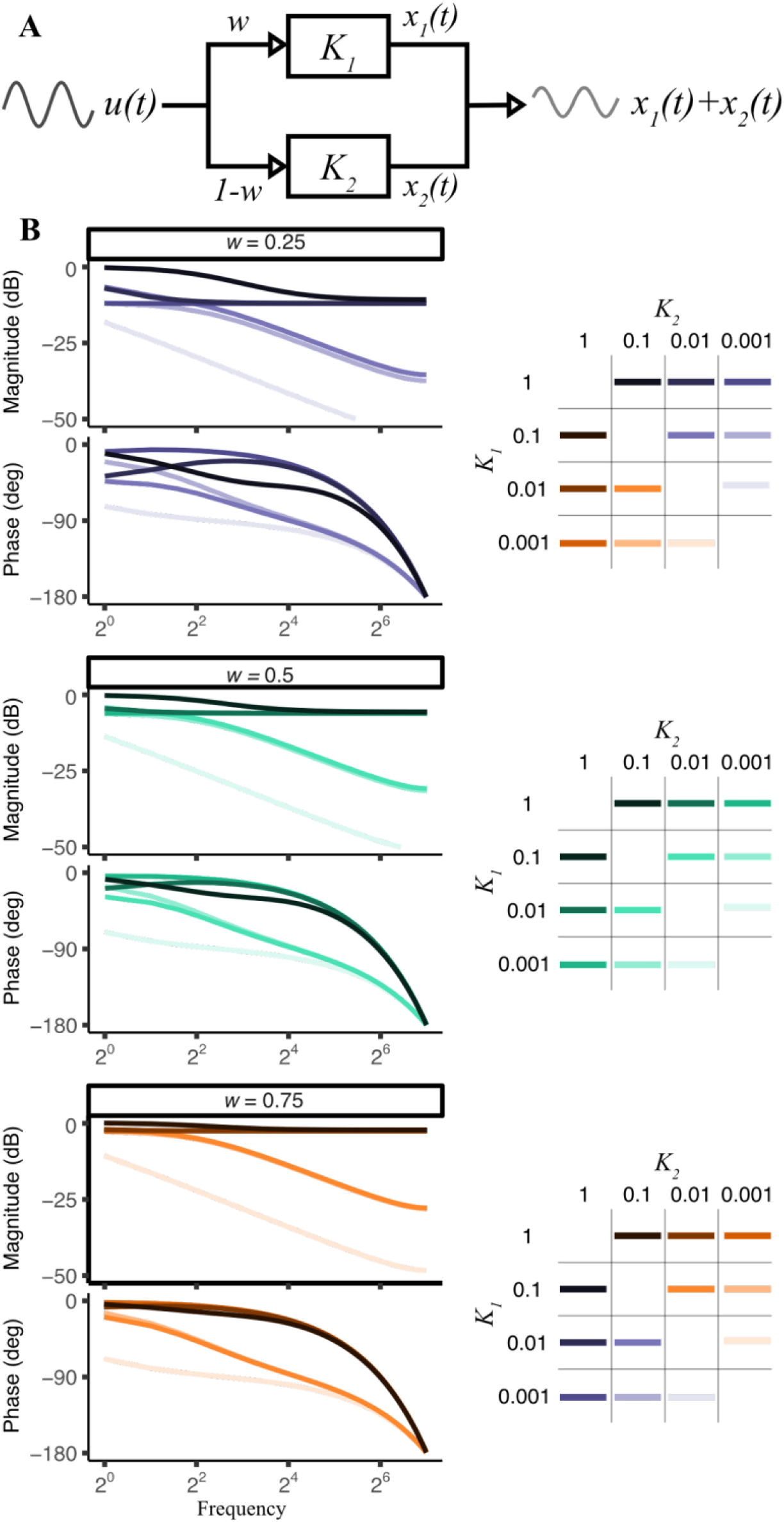
**A)** Representation of a dual-state parallel system. **B)** Simulations of the system’s frequency response, for varying values of the free parameters—namely, adaptation rates *K*_*1*_ and *K*_*2*_, and weight *w*.

In this case, the system has three independent parameters, namely the two state gains *K*_*1*_ and *K*_*2*_, and the relative weight assigned to each state *w* (Fig. 4B).

Unlike for a serial system, assigning the same weight to both states in a parallel system yields the same response of the corresponding single-state system, regardless of which weight is assigned to either state. Similarly, if the weighting parameter has the value of *w* = 1 or *w* = 0, the resulting response is that of a single-state system with gain *K*_*1*_ or *K*_*2*_, respectively. Note that the two states of the system are symmetrical: therefore, the response of a system with parameters *K*_*1*_, *K*_*2*_, *w* will be identical to that of a system with parameters *K*_*2*_, *K*_*1*_, 1–*w*, i.e. where the gain of each branch is inverted and the weight assigned to the first state is 1–the original weight (Fig. 4B).

## Materials and methods

### Experimental procedure

In order to obtain the empirical data to be used in the system identification procedure, we measured participants’ motor error in a series of rapid reaching tasks, in which they were exposed to different conditions of measurement uncertainty coupled with sinusoidal perturbations of varying frequency. Prior to that, we conducted an experiment using an alternative forced choice localization task to determine the uncertainty associated with various levels of visual blur. The results of the localization task were used to select the blur of the visual feedback in the different uncertainty conditions of the adaptation task.

#### Participants

Seven participants (4 females, 3 males, age 26.3 ± 6 years, 6/1 right/left-handed) took part in the localization task. The sample size was chosen based on a priori power analysis conducted with the G*Power application version 3.1.9.6 (41), as it would achieve 80% power for detecting a medium effect at a significance level of ɑ = 0.05. Ten participants (5 females, 5 males, age 22.1 ± 2 years, all right-handed) volunteered in the adaptation task. They were all informed of the general purpose of the experiment but were naïve with respect to the specific research questions, and received 7,50 €/h compensation or course credit for their participation. They all had normal or corrected-to-normal vision and no motor impairment. The study was approved by the ethics committee of the University of Ulm and conducted in accordance with the Declaration of Helsinki.

#### Setup

Visual stimuli were presented on a large LCD display (Sony 65X8505B, dimensions 144 × 81 cm, 1920 × 1080 pixels), placed at a distance of 85 cm from the sitting participant, in an otherwise dark room. The desk between screen and participant was customized so that on top of it was a scaffold holding a chin rest, and a black rigid cloth that prevented participants from seeing their own hand (Fig. 5A). Stimuli were programmed in Matlab R2017a and Psychtoolbox (42,43).

**Figure 5:**
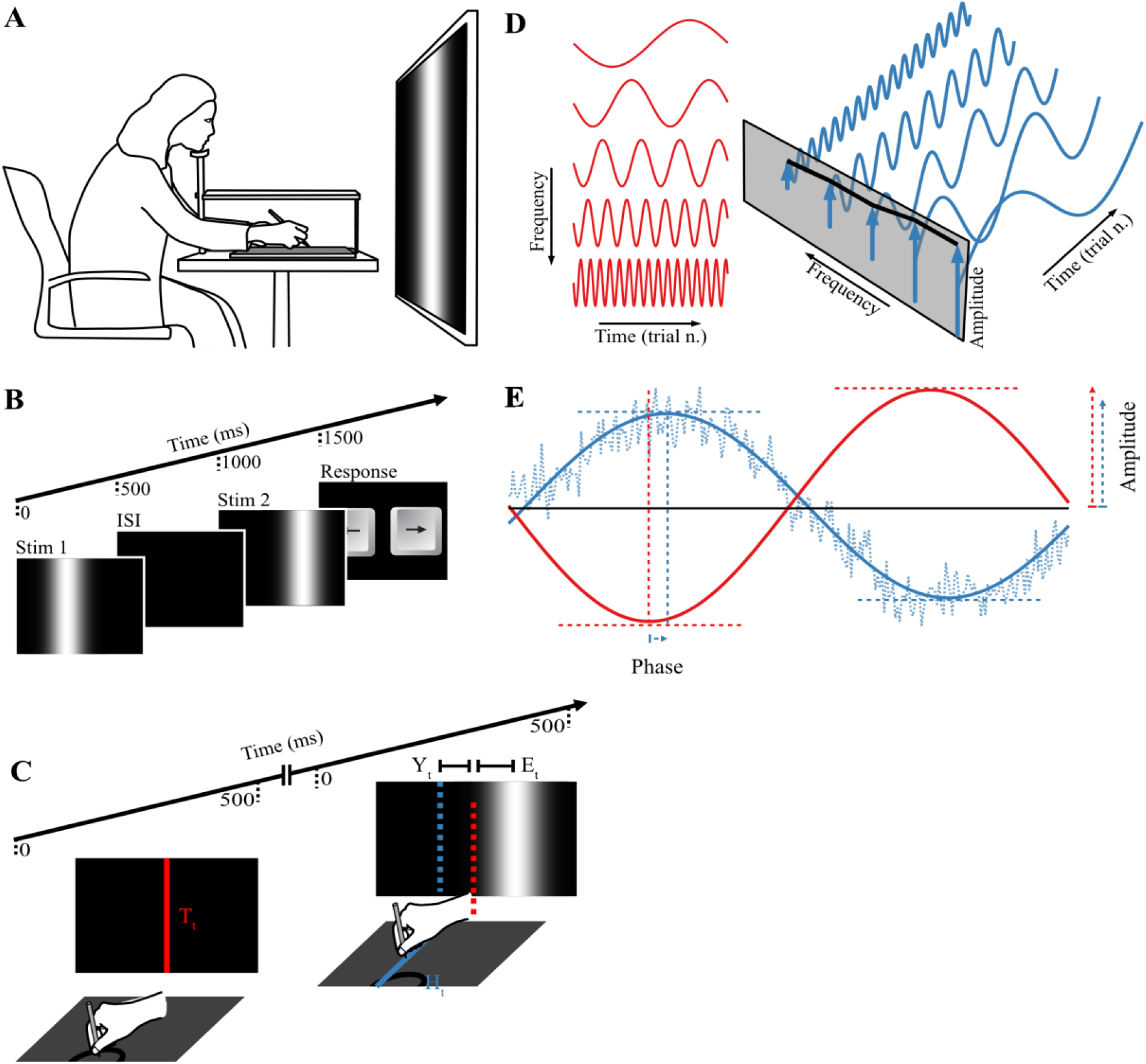
**A)** Experimental setup. **B)** Trial procedure for the localization task. **C)** Trial procedure for the adaptation task. **D)** Representation of perturbation functions (red lines) and relative adaptation responses (blue lines). Each perturbation is characterized by its frequency, i.e. the number of period repetition per run of 256 trials. The adaptation response associated with each perturbation has the same frequency, but a difference in amplitude and a phase delay. The responses associated with all tested frequencies define the frequency response, or response in the frequency domain, of the system under investigation (black line). **E)** Definition of amplitude of and phase lag of the response (blue line) with respect to the perturbation (red line).

### Localization task

Visual stimuli consisted of a vertical white line on a black background, as tall as the screen, and blurred along the horizontal direction with a Gaussian filter with varying standard deviations, starting from 0° visual angle (no blur), 8°, 12°, 16°, to 20°, resulting in 5 blur levels. The non-blurred stimulus consisted of a white vertical line of 0.5° width. Two identical stimuli were presented in rapid temporal succession (Fig. 5B; presentation time for each stimulus: 500 ms; inter-stimulus-interval: 500 ms). The first stimulus appeared at a random horizontal location on the screen within an area of width 8° centered in the middle of the screen. The second stimulus appeared on one of eight fixed distances to the left or right with respect to the first. The distance between stimuli varied linearly in a range of ±4.6° for all the blur levels except 0° where it was ±1.2°. Participants had to respond on which side they perceived the second stimulus to have appeared with respect to the first by pressing the corresponding arrows on a keyboard. Each of the eight distances of the comparison stimulus was repeated 20 times, resulting in a total of 160 forced-choice trials for each blur level condition. All trials of a given blur level were presented en bloc with a randomized order. To avoid fatigue, a non-mandatory break after every 20 trials was signaled by the appearance of a gray screen between trials.

#### Data analysis

Participants’ responses in the localization task were analyzed with a Generalized Linear Mixed Model (GLMM) to estimate the just noticeable difference (JND) associated with each blur condition (44). The model was fitted with R version 4.0.2 using the `lme4` package (45). We then calculated JND estimates and their bootstrap-based confidence intervals using the `MixedPsy` package Moscatelli et al., 2012; (46).

### Adaptation task

In each trial, a red target line (width 0.5°) appeared on the screen at a random horizontal location within an 8° wide area centered in the middle of the screen. Participants were instructed to point on a graphics tablet (WACOM Intuos3 A3; active area 48.8 cm x 30.5 cm). They had to point to the location they perceived as corresponding to that of the target on the screen as accurately as possible (Fig. 5C). That is, their task was to minimize the distance between target and feedback position (i.e. the feedback error *E*_*t*_, Eq. 5). The pointing movement was done with a stylus held in their preferred hand. The feedback stimulus consisted of a white vertical line blurred along the horizontal dimension, identical to the one described for the localization task. Only a subset of the blurs tested in the localization experiment was used in the adaptation experiment, namely: 0° (no blur), 12° (intermediate blur) and 20° (large blur).

The experiment included separate runs consisting of 256 trials each. The mapping between tablet and screen coordinate changed on every trial within a run along the horizontal direction, according to a sinusoidal perturbation function in the form:

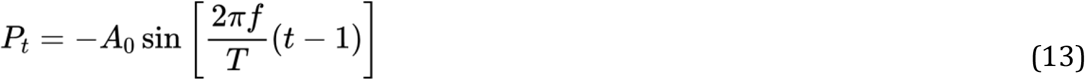

where *A*_*0*_ was the perturbation amplitude set to 3°, *T* was the total number of trials within a run (*T* = 256), and *f* was the frequency of the sinusoidal perturbation, corresponding to the number of times that the full period of the sinusoidal function was repeated within a run. Following the method proposed by Hudson and Landy (38), the negative sign in Eq. (13) was chosen out of convenience for subsequent analysis, so that the adaptation response would follow a positive sinusoidal function (Fig. 5E). Also, a lag of 1 trial was introduced so that the first trial of each run would have no perturbation. We presented 7 sinusoidal frequencies, chosen as integer powers of 2 and ranging between 2^0^ = 1 and 2^6^ = 64 cycles/run (Fig. 5D). We tested every sinusoidal frequency with each of the three blur levels (i.e. 3 blurs x 7 frequencies), resulting in 21 runs of 256 trials. All 7 frequencies of a given blur level were tested en bloc in randomized order, with each blur level tested on a different day. At the beginning of each testing day participants performed 100 training trials of the reaching task without any perturbation, but with a feedback blur consistent with that of the rest of the day. The order in which the blur levels were tested across days was counterbalanced between participants.

Every reaching movement began in a small area at the center/bottom of the tablet, delimited by a plastic half-circle ridge of diameter 3.5 cm (Fig. 5C). In order to trigger the presentation of a new target, participants pressed a button on the grip of the stylus while holding its tip inside the ridged half-circle (starting area). They were allowed to keep their non-preferred hand on the ridge in order to find the starting area more easily. While participants kept the stylus within the starting area, the target was visible on screen for a maximum of 500 ms. Once the stylus left the starting position, participants needed to make a fast reaching movement towards the upper half of the tablet. The endpoint position where the stylus first touched the tablet counted as the response; however, it did not count as valid unless the stylus moved at least 9 cm away from the starting position along the sagittal plane. If the position of the stylus passed this limit, a feedback stimulus appeared and remained on the screen for 500 ms. Otherwise, no feedback appeared and the trial had to be repeated.

The tablet’s coordinates mapped univocally to the coordinates of the screen, with the tablet’s boundaries corresponding to those of the screen. Although the devices did not share the same dimensions, no participants demonstrated difficulties in understanding the applied scaling.

#### Data analysis

Since no perturbation occurred along the vertical axis of the screen, we did not analyze data relative to the vertical endpoint coordinate of the reaching movement. To evaluate the adaptation signal in response to a sinusoidal perturbation, we considered the motor error *Y*_*t*_, defined as the difference between the target coordinate and the horizontal coordinate of the reaching endpoint position in the unperturbed mapping. We assumed that the adaptation response to a sinusoidal perturbation with frequency *f* would have the same frequency but a smaller amplitude and a delay, i.e. a phase difference with respect to the perturbation (Fig. 5D-E). The adaptation response was modeled accordingly (38):

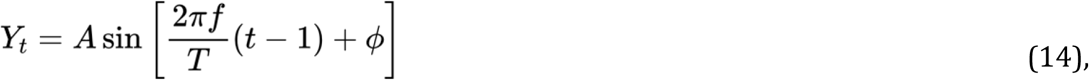

where *A* is the amplitude (0 ≤ *A* ≤ *A*_*0*_) and *φ* is the phase of the adaptation response (0 ≤ *φ* < 2π).

Eq. (14) can be rewritten in analogous terms by the properties of trigonometric functions as follows:

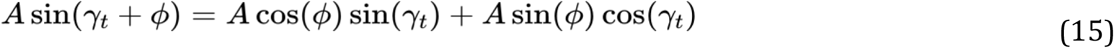

with 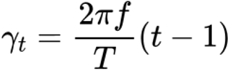.

The identity in Eq. (15) allows to quantify the amplitude and phase of the response by fitting the adaptation response with a linear mixed model (LMM) in the following form, for each combination of frequency and visual blur:

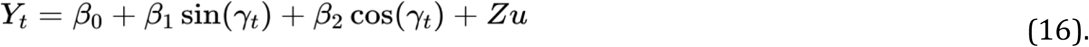

Where *β*_*0*_, *β*_*1*_, *β*_*2*_ represent the fixed-effect parameters of the model, and *Zu* accounts for the random effects. Of the fixed-effect parameters, *β*_*0*_ describes the constant response bias associated with the response, and is expected to vary randomly for each participant (38). In line with Eq. (15), parameters *β*_*1*_ and *β*_*2*_ can be rewritten as:

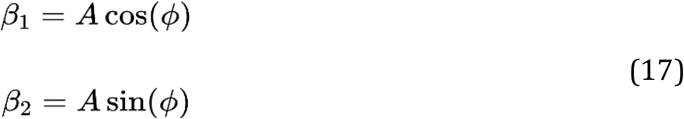

In this way, we can extract amplitude and phase parameters (*A* and *φ* in Eq. 14) given the linear predictors *β*_*1*_ and *β*_*2*_, by arranging and combining them in light of basic trigonometric identities. For the amplitude, we have:

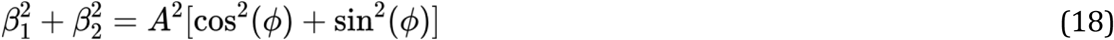

which yields

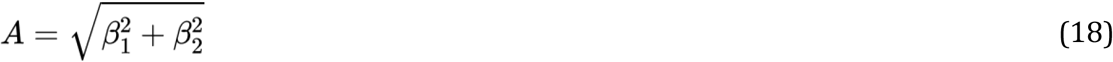

given the Pythagorean identity: cos^2^(*ϕ*) + sin^2^(*ϕ*) = 1.

For the phase shift, by considering the definition of the angle’s tangent: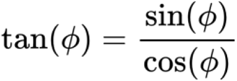, we obtain:

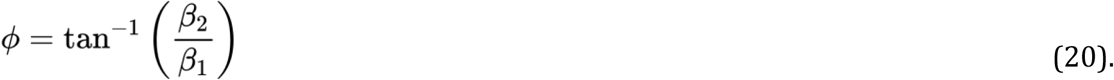

The intercept parameter *β*_*0*_ represents a constant bias in the response. Since such constant bias, which is small and variable across participants, does not affect the frequency response or the amplitude of adaptation, it was removed from the data and excluded from further analysis, without affecting the other model parameters.

LMMs for each blur and frequency (21 models in total) were fitted in R with the `lme4` package. Amplitude and phase parameters for individual participants were extracted from the combination of fixed and random effects of each coefficient of the LMMs. We calculated Bootstrap-based confidence intervals for estimating parameters at the population level using the `MixedPsy` package. Finally, the amplitude of the response (both at the individual and sample level) was transformed in magnitude, i.e. it was normalized with respect to the amplitude of the input perturbation and expressed in decibel (dB): 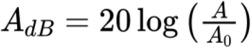.

Magnitude and phase fully describe the adaptation response to a given sinusoidal perturbation in the frequency domain (39). Taken together, these parameters provided the frequency response of adaptation in the frequency range of the presented perturbations for each level of feedback uncertainty (Fig. 5D).

### System Identification

Given the frequency responses extracted from the adaptation signals, we used the state-space parametrizations for the model architectures described in the previous section to estimate the corresponding best fitting models. The state-space model estimation was performed in Matlab R2017a using functions from the System Identification Toolbox (40). In particular, we used the function `greyest`, which performs a grey-box estimation of a LTI system given the frequency response of the empirical data and the specific relations between system parameters.

For each participant and condition of measurement uncertainty, we estimated the following possible models from the associated frequency response:

- a one-state system (Eq. 6) characterized by the single adaptation rate *K*;
- a dual-state system with serial architecture (Eq. 11), described by two adaptation rates *K*_*1*_ and *K*_*2*_;
- a dual-state system with parallel architecture (Eq. 12), described by the adaptation rates *K*_*1*_ and *K*_*2*_, and the relative weight *w* assigned to each state.

Without loss of generality, in the dual-state configurations, we always assigned *K*_*1*_ to the faster, and *K*_*2*_ to the slower rate. Accordingly, for the system with a parallel architecture, the weighting factor *w* refers to the weight assigned to the state with faster rate and *(1-w)* to the state with the slower rate.

In the system identification function, we constrained every parameter to be limited between 10^−15^ and 1. The small bias added to the lower threshold was included to avoid undefined results in subsequent analyses in case parameters were estimated at exactly 0.

In order to evaluate which of the three considered systems was best at fitting the adaptation response, we used the Bayesian information criterion (BIC) to compare the models. For each frequency response we selected the model with the lowest BIC score and evaluated the goodness of fit relative to the other models in terms of the difference between BIC values (ΔBIC). Models with ΔBIC between 0 and 2 were not considered different from each other; models with ΔBIC between 3 and 10 were classified as little though still significantly different; models with ΔBIC greater than 10 were considered strongly different from each other (47,48).

### Effect of uncertainty on identified parameters

Once we identified the parameters of the three system architectures, we evaluated how the JND associated with the visual blur of the feedback (i.e. their measurement uncertainty) affected the parameter values. Since the system with two states in parallel provided the best fit of the three, we selected this model for further considerations. In general, we expected that the system, regardless of its structure, would adapt slower as an effect of the increase in feedback uncertainty. In case of a dual-state system with a parallel architecture, this could be achieved in two ways: (1) by decreasing one—or both—adaptation rates, and/or (2) by assigning more weight to the state with the lower adaptation rate. We fitted LMMs to adaptation rates and relative weight in order to test our hypotheses.

#### Adaptation rates

We expected that adaptation rates, if they changed, would become smaller as the uncertainty associated with visual feedback increased. We modeled this relationship with an exponential decay in the form:

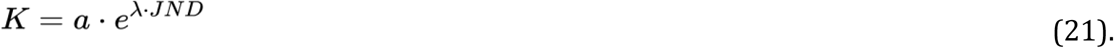

We applied a logarithmic transformation to both elements in Eq. 21 in order to fit a LMM as follows:

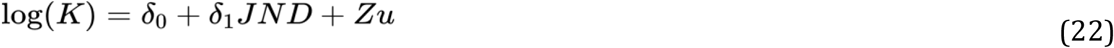

where *δ*_*0*_ = log(*a*) and *δ*_*1*_ = *λ*.

The model in Eq. 22 was applied to both rates, *K*_*1*_ and *K*_*2*_ of the systems. Significance of the fixed-effect parameters was set at 0.05 and evaluated with likelihood ratio (LR) tests.

#### Relative weight

We expected the weight assigned to the faster state *w* to decrease as a function of feedback JND. As a first approximation, we evaluated whether such decrease could be described as a linear function with a negative slope in the form:

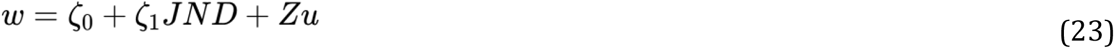

Since the weight is bound to change between 0 and 1, the model in Eq. (13) is not appropriate to fully describe the relationship between weight and feedback JND, hence it must be considered as a local approximation for the considered JND interval.

## Results

### Localization task

In the localization task, the JND increased monotonically and non-linearly as a function of the standard deviation of the gaussian blur applied in the different conditions (Fig. 6). In other words, an increase in blur of the visual feedback was associated with an increase of measurement uncertainty. To provide visual feedback in the adaptation task, we chose the three values of visual blur (0, 12, & 20°) for which the confidence intervals of the JND estimates were not overlapping (Table S1 in Supplementary Materials).

**Figure 6:**
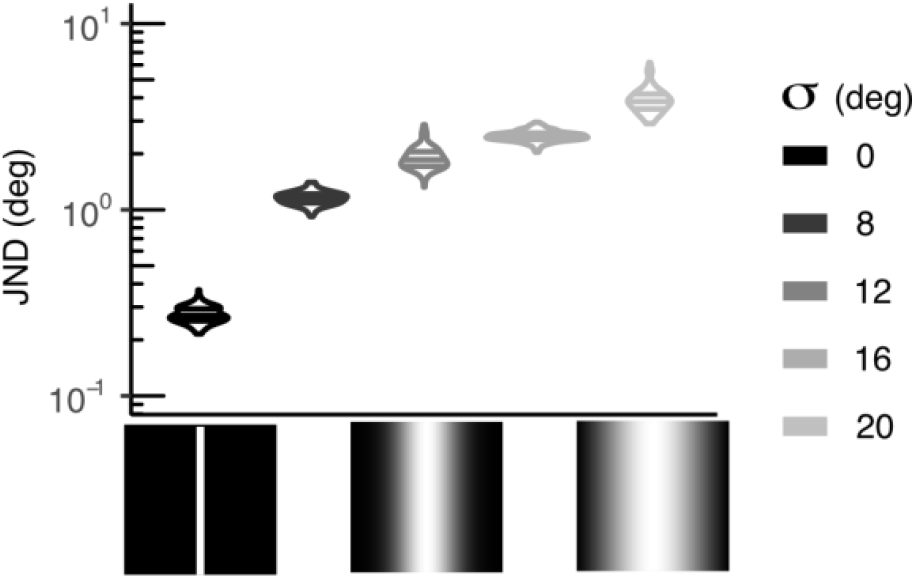
JND associated with all the blurred stimuli presented in the localization task.

### Adaptation task

The model in Eq. (16) was applied to the motor response *Y*_*t*_ in each condition of perturbation frequency and measurement uncertainty. Fig. 7 shows the sinusoidal perturbation together with averaged response across participants. As expected, the adaptation response generally followed a sinusoidal pattern with the same frequency, but a reduction in amplitude and a phase delay with respect to the imposed perturbation. The summary of magnitude and phase values of the adaptation response as a function of frequency is shown in Fig. 8A in a Bode plot.

**Figure 7:**
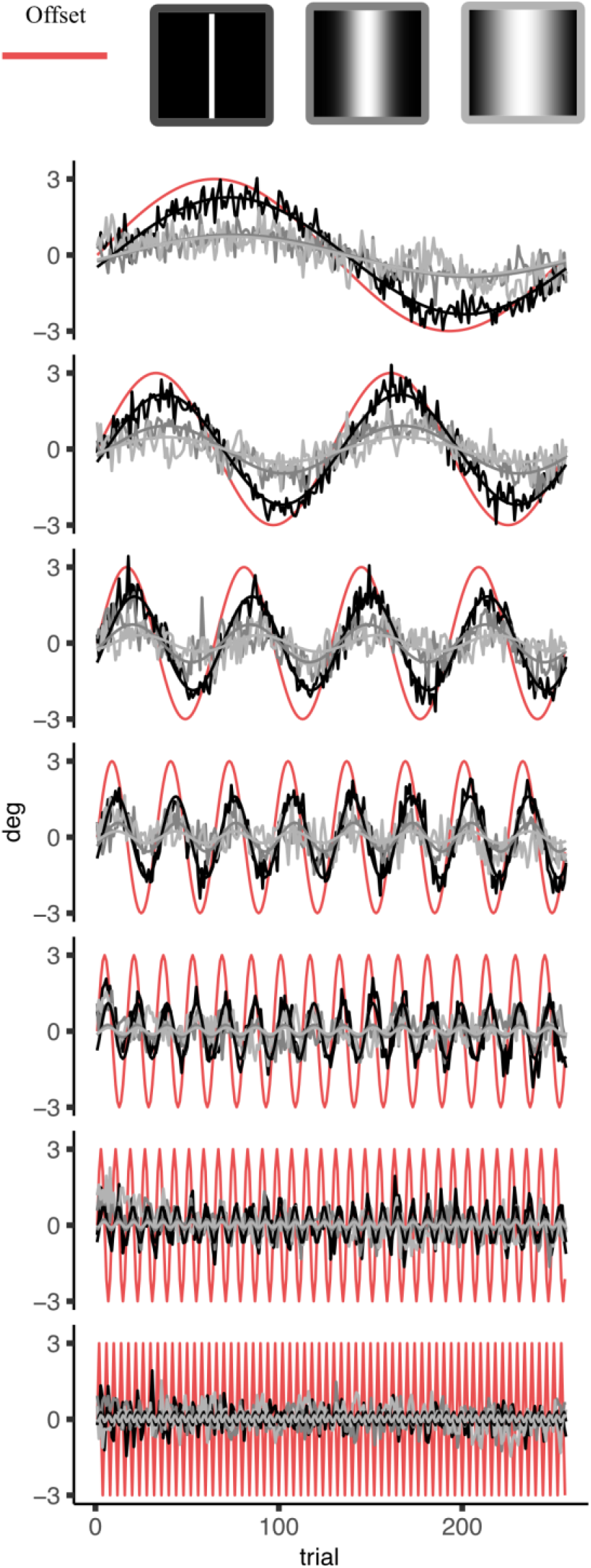
Sinusoidal fit and adaptation response averaged across participants (n = 10). The offset perturbation for each frequency condition is shown in red. Amplitude and phase of the sinusoidal curves in gray are calculated from to the fixed-effect parameters of the model in Eq. (16) through the identities in Eq. (17). In this representation, perturbations are inverted to allow the reader to visually evaluate the difference between input and response, by comparing the peaks of the two signals.

**Figure 8:**
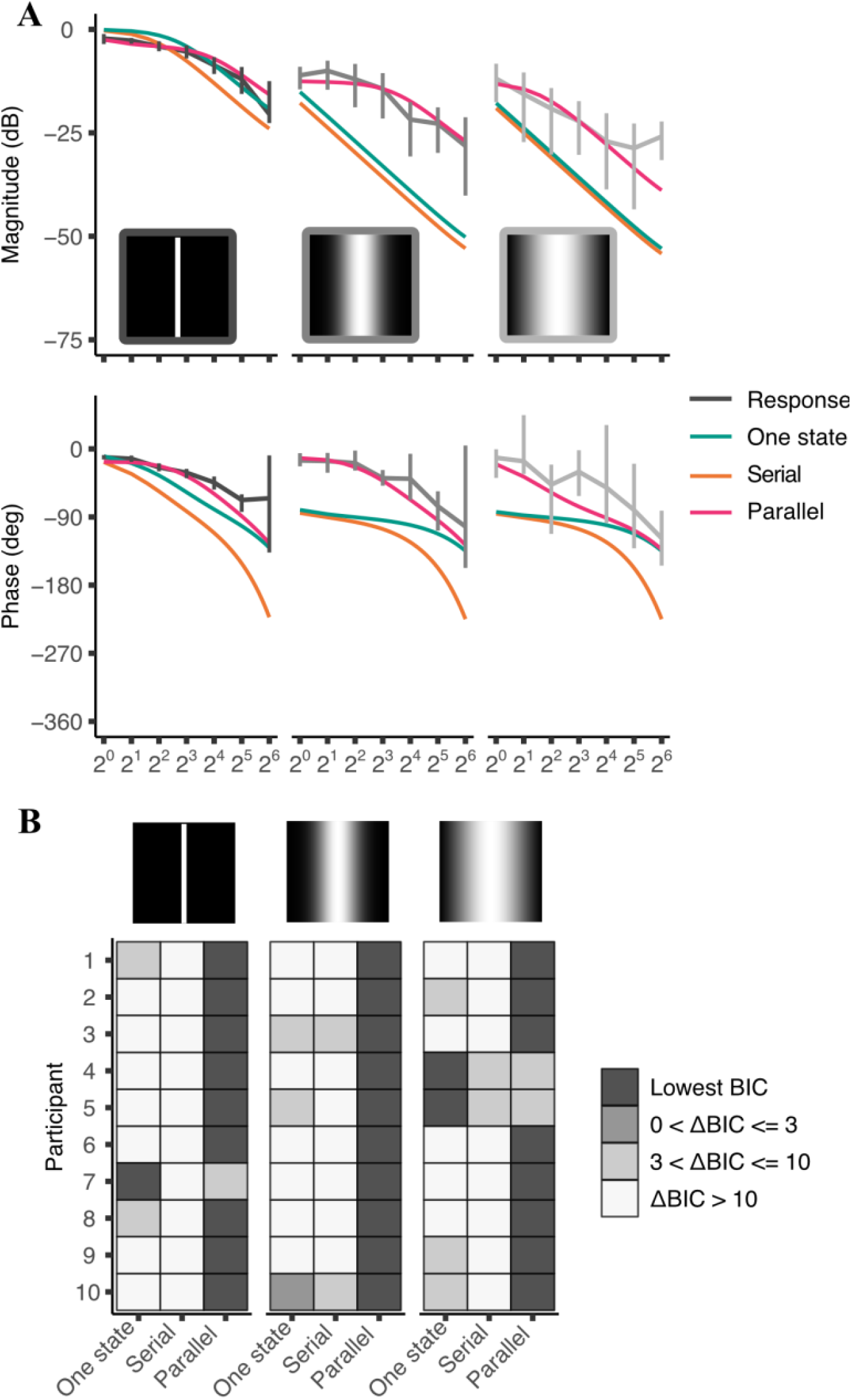
**A)** Bode plot: Comparison between frequency response for the population sample (estimate and bootstrap-based confidence intervals) and response associated with median identified parameters for the three system architectures under consideration. **B)** Model comparison according to the BIC criterion for individual fittings.

A first qualitative observation from visual inspection showed that the amplitude of the adaptation response was smaller for higher frequencies and increasing measurement uncertainty, while the phase lag increased for higher frequencies but tended to be similar across different conditions of uncertainty.

### System identification

The parameters resulting from the estimation of the three system architectures (i.e. a single state, two states in series, and two states in parallel) are summarized in Table 1.

**Table 1:** Parameters of the best fitting systems (Median and IQR) for each condition of visual blur σ.

| $\sigma$ (deg) | One-state | Serial | | Parallel | | |
| --- | --- | --- | --- | --- | --- | --- |
| | K | $K_1$ | $K_2$ | $K_1$ | $K_2$ | w |
| 0 | 0.145 (0.066) | 1.00 (0.605) | 0.101 (0.200) | 0.313 (0.142) | 0.015 (0.014) | 0.616 (0.214) |
| 12 | 0.005 (0.007) | 1.00 (0.00) | 0.00 (0.00) | 0.226 (0.094) | 0.001 (0.002) | 0.234 (0.141) |
| 20 | 0.003 (0.005) | 1.00 (0.00) | 0.00 (0.00) | 0.073 (0.105) | 0.00 (0.001) | 0.233 (0.285) |

As is apparent from Fig. 8A, the system with a parallel architecture was the most effective in capturing the characteristics of the frequency response for all conditions of measurement uncertainty. Accordingly, it had the lowest BIC score 27 out of 30 times (Fig. 8B). In four cases, the single-state system was indistinguishable from the system in parallel or provided a better fit, with a small but significant difference (ΔBIC between 3 and 10). The system with a serial architecture was the one consistently providing the worst fit, and accordingly, the one with the largest BIC scores in all cases.

### Effect of uncertainty on identified parameters

After having identified the free parameters for each of the three considered system architectures, and having established that the parallel system was the best amongst the candidate models, we focused on the relationship between measurement uncertainty (i.e. the feedback JND) and the identified parameters only for this parallel model.

#### Adaptation rates

The LMM fitted to the fast learning rate *K*_*1*_ (Eq. (22); Fig. 9A, black symbols) revealed an intercept of 0.334 (95% CI: [0.259, 0.43]) and a decay constant λ = −0.297 (95% CI: [−0.496, −0.096]) indicating that the fast adaptation rate became slower with increasing blur. The fixed effect accounting for the decay of the fast adaptation rate as a function of JND was significant (χ_1_ = 6.922, p = 0.009). For *K*_*2*_ (Fig. 9A, gray symbols), the intercept was very close to 0 (∼ 10^−4^, 95% CI: [10^−7^, 1.361]). Still the decay constant, albeit small, was significantly different from 0 (λ = −4.66, 95% CI: [−7.901, −1.284]; χ_1_ = 7.334, p = 0.007).

**Figure 9:**
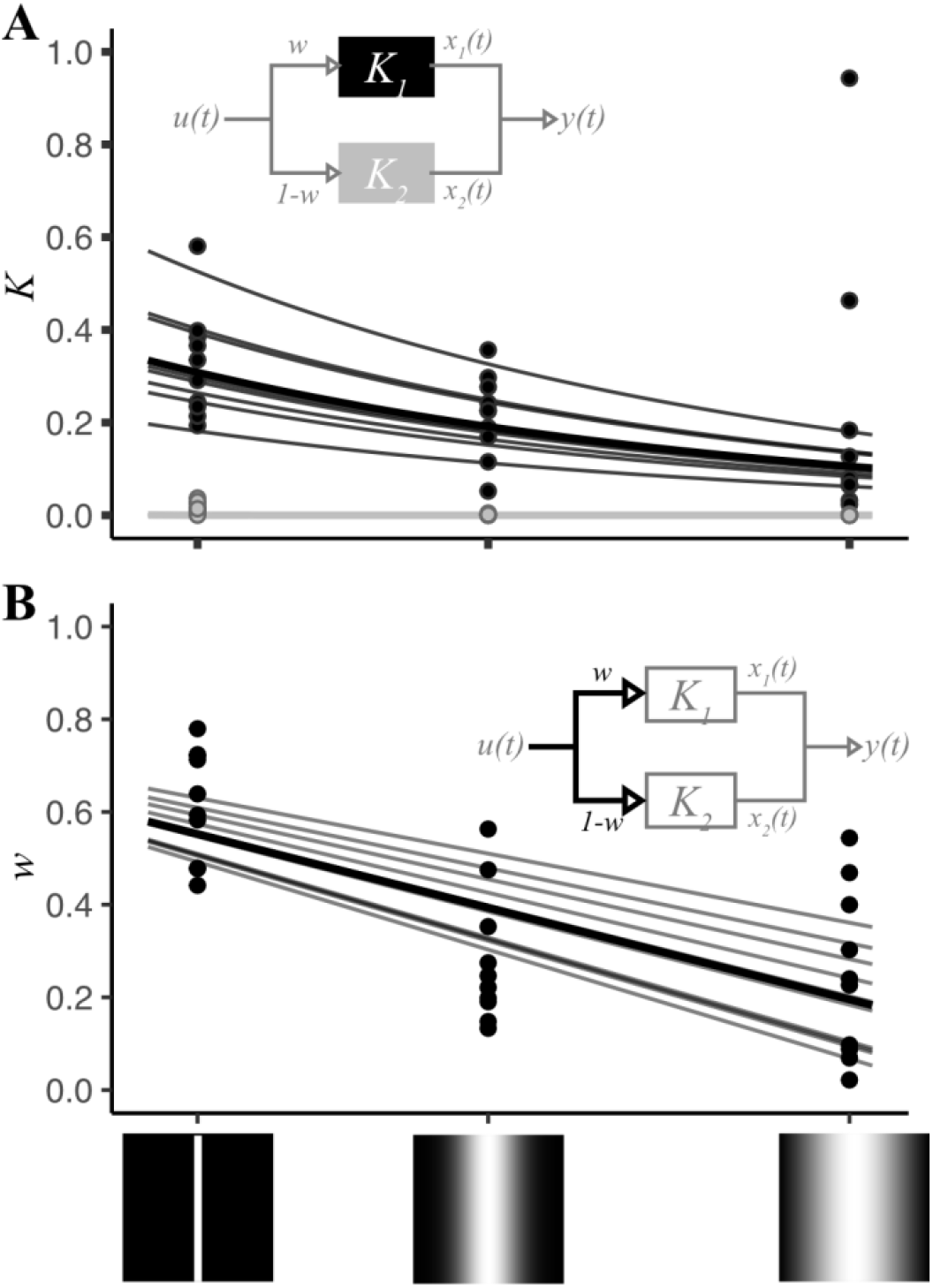
**A)** Values and fitted models of learning rate parameters *K*_*1*_ and *K*_*2*_ for the system with two states in parallel, as a function of feedback JND. **B)** Weight (values and fitted model) assigned to the fast learning state (*K*_*1*_) in the parallel system as a function of feedback JND.

#### Weighting factor

For parameter *w* (Fig. 9B) the model in Eq. (23) had a negative slope (ζ_1_ = -0.099, 95% CI: [−0.135, −0.062]. χ_1_ = 16.709, p < 0.001) indicating that the weight assigned to the faster state decreases with increasing uncertainty (i.e. the slower state was weighted more as feedback JND increased).

## Discussion

We investigated visuomotor adaptation of pointing movements under different conditions of sinusoidal perturbation and feedback uncertainty, and applied system identification techniques to determine which system architecture provided the best fit to the empirical data.

Our results confirm and extend those of previous studies: Adaptation responses were successfully fitted by sinusoidal functions having the same frequency, but different amplitude and phase with respect to the perturbation (38). This indicates that participants were able to adapt to sinusoidal offsets at various speeds. In particular, the response had higher amplitude and smaller delay at lower frequencies and tended to degrade for higher frequencies, in accordance with the prediction that adaptation to systematic perturbations resembles the response of a passive low-pass filter. Moreover, adaptation performance was affected by the modulation of the feedback’s visual blur, with higher uncertainty (i.e. larger blur) associated with less complete, slower adaptation (10).

When applying system identification procedures to the adaptation response expressed in the frequency domain, we found that a dual system with a parallel architecture provided the best fit to our data for all levels of feedback uncertainty. The parallel architecture provided a better fit than the dual-state system with a serial architecture 100% of the time. 90% of the time the parallel architecture was better than a single-state system taking also model complexity into account. In the remaining 10% of cases in which the single-state system outperformed the parallel system, the best fitting parameters were such that the two systems were hardly distinguishable: namely, the assigned weight in the parallel system architecture was close to either 0 or 1. Such result indicates that in the majority of cases the system identification algorithm converged to a solution well distinguishable from that of a single-state system: Namely, the learning rate of the two independent states differed from each other, and both states were assigned a non-zero contribution.

As an effect of increasing feedback uncertainty, the values of all three identified parameters (i.e. the two adaptation rates *K*_*1*_, *K*_*2*_, and the weight assigned to the faster state *w*) decreased significantly, even though the adaptation rate of the slower state *K*_*2*_ was close to zero in all uncertainty conditions (Table 1, Fig. 10). Nevertheless, not only was the weight assigned to its contribution (i.e. *1–w*) significantly different from zero in all conditions, but it also increased for larger feedback blurs.

**Figure 10:**
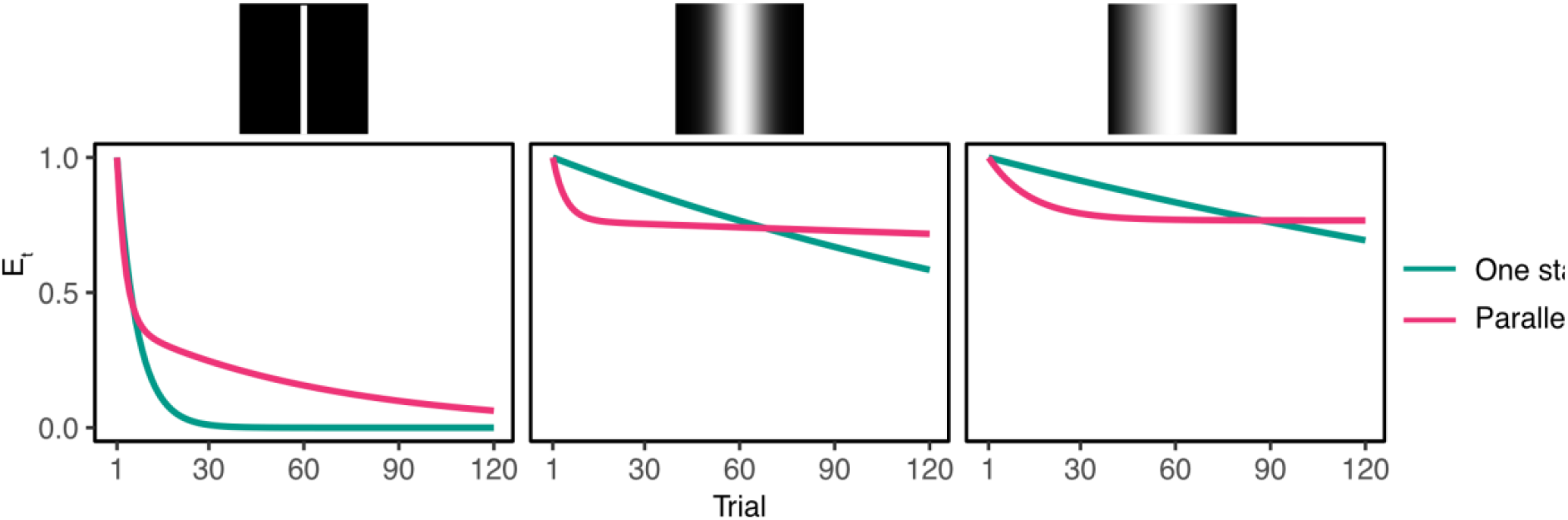
Simulations of the impulse response to a constant perturbation under different conditions of feedback uncertainty. The simulations are relative to the systems with one state (green lines) and two states in parallel (magenta lines) identified from the frequency responses obtained in the sinusoidal adaptation experiment.

To better understand the implications of this aspect of the identified model, let us consider the simulations of the response to a step offset associated with both the single-state and the parallel system (Fig. 10). For a single-state system (Fig. 10, green lines), the error decreases according to an exponential decay function, whose time constant is defined by the value of the free parameter *K*. Since the identified values of *K* under medium and high uncertainty tend to zero (Table 1, Fig. 1S), the rate of decay of the error is very slow in these two conditions, and can be even approximated by a linear decrease in the displayed interval of 120 trials (Fig. 10, middle and right panel). Regardless, at the end of the exponential decay, error is fully eliminated and performance returns to baseline levels in all cases (given enough trials), as is predicted by the Kalman filter model. The response of a dual-state parallel system (Fig. 10, purple lines) differs qualitatively from that of a single Kalman filter. For the condition under lower uncertainty (Fig. 10, left panel), the error does tend to revert to baseline, although after a large number of trials, and with a decay profile dependent not only on the faster rate, but also on the slower rate as well as their relative weight. Yet, since the slow adaptation rate *K*_*2*_ is several times smaller than the fast adaptation rate *K*_*1*_ (Table 1), the latter dominates for the larger part of the adaptation response in the considered interval. In conditions of medium and high uncertainty, the value of the slow gain *K*_*2*_ is even smaller, to the point that it can be approximated to zero without a significant decrement in the fitting accuracy. As a result, the rate of the error’s exponential decay is defined by the value of the fast gain *K*_*1*_, while the relative weight w determines the asymptote of the decay. Notably, this prediction of the parallel dual-system is highly consistent with the empirical results obtained by Burge and colleagues (10). In their study, results showed the presence of an incomplete compensation in the adaptation response, i.e. an asymptote of the exponential error decay larger than zero, which increased in conditions of higher uncertainty. Such residual error bias was nevertheless excluded from the proposed model, because it did not appear to affect the estimate of the adaptation rate (10,24). The presence of a constant residual error has been described in several studies on visuomotor adaptation e.g. (10,31,32,49–51), in full disaccord with the predictions of the Kalman filter model.

Instead, state-space models can account for incomplete offset compensation, by introducing a forgetting factor which describes the tendency for the system to return to a baseline, unperturbed state on every trial (32). In this framework, incomplete adaptation is interpreted as the effect of the balancing between multiple retention and learning processes (described by parameter A and B in the state estimation update equation, respectively; cf. Eq. 7) operating at different rates (31). In their study, van der Kooij and colleagues (31) proposed that this phenomenon is not a fallibility of the adaptation system, instead it is a mechanism by which the system can react in a conservative fashion to possibly transient changes in the environment (e.g. unstable external perturbations or fatigue) while being able to adjust effectively to more permanent changes, such as body modifications following development. Nevertheless, it is unclear from modeling studies within the state-space framework how such a combination of learning and retention factors are actually implemented in the physical plant of the system responsible for adaptation. Our results suggest that incomplete adaptation is the result of two concurrent mechanisms: On the one hand, the states of the system contributing to adaptation in parallel respond to the perturbation with different rates, which are sensitive to the change in the environmental statistics and therefore decrease their learning rate in presence of higher feedback uncertainty, thus slowing down the overall adaptation; On the other hand, the contribution to adaptation assigned to either state also changes, so that with increasing uncertainty, the response of slower state determines the larger part of the system’s response. As the learning rate of the slow, dominating state tends to zero, this translates into little or no adaptation once the contribution of the faster state is extinguished, and therefore to a residual error bias. It is likely that the weight assigned to each state of the system, just like the learning rates assigned to the states themselves, is optimally tuned to the statistics of the environment. However, further research is needed to investigate the statistical determinants of contribution assignment in adaptation.

By showing evidence supporting the hypothesis that multiple estimators act like optimal observers, each one contributing to the adaptation response by processing the input information in parallel, we can provide a better insight on the mechanisms of sensorimotor adaptation, as well as the way they are physically implemented. On every trial, a motor plan is actuated based on the mapping estimated from error in previous trials, and a new error is perceived in the form of a visual displacement with respect to the intended target. Given this error signal, the controller must establish which one of the systems involved is more likely to have caused the perturbation. On the one hand, the error can be attributed to an incorrect action plan performed by the motor system; on the other hand, the cause of error can be imputed to a misperception of the input signal by the visual system. Following error assignment, the controller must establish which part of the system is responsible for correcting the perturbation error in subsequent trials, in order to update the state estimation with one more appropriate for the current environmental conditions. In scenarios of error reduction tasks under ambiguous, redundant conditions, the controller typically assigns error reduction to the component that is considered as more likely to have caused the error (52). In their study, White and Diedrichsen (52) showed that, in bimanual motor tasks, right-handed participants corrected the perceived error induced by a perturbation more with their left hand, even if the right hand was generally able to correct for error faster in non-redundant tasks. In analogy with this phenomenon, in visuomotor adaptation tasks like the one in our study, the decision regarding which estimator is more likely to have caused the error should result in the appropriate contribution to error reduction, i.e. the selection of weight attributed to the different states of the system. Our results show that, under conditions of higher uncertainty, the controller assigns the task of error reduction to the state known to be slower in learning. This finding is in line with the idea that error correction is performed by the state that is more likely to have caused the error in the first place.

It is likely that reducing the entire plant of sensorimotor adaptation to two distinct systems is an oversimplification (21,33). By extending the model proposed here, we can test increasingly complex implementations of the adaptaing system that takes into account the constraints imposed by the physical plant of the sensorimotor system. For example, we can test a more articulated implementation of the system by adding other states to the parallel system, either in series or in parallel. Such configurations would account for the contribution of different sensory and motor components, together with the internal complexity of each individual component. Since such an approach would result in a rapid increase of the free parameters of the model, with the risk of incurring in model overfitting with our current dataset, we limited our investigation to systems including only two independent states.

In conclusion, our results show that the adaptation response to visuomotor perturbations is well approximated by a linear time-invariant system with two states in parallel. Unlike the more commonly used model with one single Kalman filter, such an architecture captures an often described characteristic of adaptation, namely incomplete error compensation in response to a constant offset, which results in a nearly constant residual bias. Under conditions of increasing feedback uncertainty, the learning rate of both states slows down, while the weight assigned to the slower state increases, suggesting that the controller assigns a larger contribution in error reduction to the system which is more likely to have caused the error. Such an explanation is consistent with the scenario in which the motor system adapts quickly, while the visual system adapts slowly.

## Acknowledgments

We thank Lynn Matits for assistance in data collection, and Chris Geekie for suggestions on language and style of the manuscript. Funded by the Deutsche Forschungsgemeinschaft (DFG, German Research Foundation) – Projektnummer 251654672 – TRR 161.

## Supplementary material

**Table S1:** Estimate and confidence intervals of JND in the localization experiment, in degrees of visual angle. Rows with gray background are those used in the adaptation task.

| Blur | JND Estimate [95% CI] |
| --- | --- |
| 0° | 0.273 [0.220, 0.334] |
| 8° | 1.159 [0.968, 1.393] |
| 12° | 1.878 [1.494, 2.650] |
| 16° | 2.479 [2.139, 2.846] |
| 20° | 3.874 [2.937, 5.634] |

**Figure 1S:**
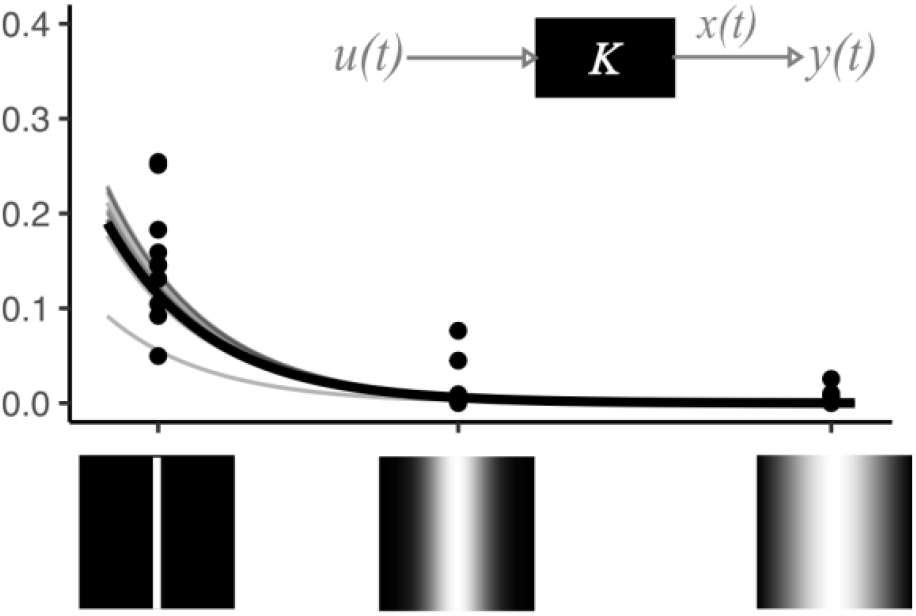
*Single-state system:* For the single-state and the parallel system, learning rates followed an exponential decay as a function of the JND of the visual feedback. The learning rate (Fig. S1) had a starting value of 0.19 (95% CI: [0.003, 3.319]) and decreased with a decay constant λ = −1.846 (95% CI: [−2.913, −0.355]). The fixed effect accounting for the decay of the learning rate was statistically significant (χ_1_ = 6.724, p = 0.01).

